# Genetic epidemiology of human neutrophil antigen variants suggest significant global variability

**DOI:** 10.1101/2022.04.28.489826

**Authors:** Mercy Rophina, Vinod Scaria

## Abstract

Human neutrophil antigens possess significant clinical implications especially in the fields of transfusion and transplantation medicine. Efforts to estimate the prevalence of genetic variations underpinning the antigenic expression are emerging. However, there lacks a precise capture of the global frequency profiles. Our article emphasizes the potential utility of maintaining an organized online repository of evidence on neutrophil antigen associated genetic variants from published literature and reports. This in our opinion, is an emerging area and would significantly benefit from the awareness and understanding of population-level diversities.

## Introduction

Granulocytes constitute the major proportion of human leukocytes and are an inevitable part of the innate immune system. As a significant majority (∼99%) of the granulocyte population primarily comprises of neutrophils, a family of glycoprotein epitopes expressed on granulocytes have been collectively named as the human neutrophil antigens (HNAs) (Browne et al., 2021),(Flesch & Reil, 2018). Based on the serologically defined epitopes, these antigens are classified into five major groups/systems namely HNA-1 to HNA-5. Molecular characterization revealed that antigens of HNA class 1-5 are encoded by FCGR3B, CD177, CLT2, ITGAM and ITGAL genes respectively (Flesch & Reil, 2018). The FCGR3B gene encoding alleles of the HNA-1 system spans chromosome 1q23-24 and also carries a high degree of homology with the FCGR3A gene (Ravetch & Kinet, 1991), (Ravetch & Perussia, 1989). CD177 glycoprotein is produced by the CD177 gene located on chromosome 19q13.2 whereas the molecular functions of CD177P, a pseudogene located downstream of CD177 still remains unclear (Bettinotti et al., 2002), (Caruccio et al., 2006). Recently, a decade ago, the HNA-3 antigen was found localized to CTL2, encoded by SLC44A2 gene spanning chromosome 19p13.1 (Greinacher et al., 2010), (Curtis et al., 2010). HNA-4a was formerly termed “Mart^a^” and is found to be expressed on the CD11b/α_M_ subunit of the α_M_β_2_ - integrin (Simsek et al., 1996). Similarly, HNA-5a antigen which was previously described as “Onda” is expressed on the α_L_ subunit of the α_L_β_2_ - integrin (Décary et al., 1979). Although HNA-1 and HNA-2 have been found specifically expressed on the neutrophil surfaces (Lalezari, 2017), classes HNA-3 to HNA-5 have a broad range of expression in other blood cells (Flesch et al., 2016).

Antibodies against HNA (auto-, iso- and allo-antibodies) have been found to possess significant clinical relevance especially in terms of transfusion and transplantation complications (Jug et al., 2021),(Key et al., 2018). Clinical disorders including alloimmune neonatal neutropenia and autoimmune neutropenia are caused by neutrophil specific antibodies (Porcelijn & de Haas, 2018),(Fung et al., 2003). In addition, almost all classes of HNA antigens (HNA-1a, 1b, 1c, 2, 3a, 4a and 5a) have been found to have significant implications in transfusion related acute lung injury (TRALI) cases (Vlaar, 2012), (Moritz et al., 2009), (Bayat & Sachs, 2012), (Caudrillier & Looney, 2012)

Limited studies have previously estimated the population frequencies of HNA antibodies(J. L. Gottschall et al., 2011),(J. Gottschall et al., 2013),(Nguyen et al., 2009),(Esmaeili et al., 2022). Various PCR-based HNA genotyping assays have been developed and continue to be the gold standard method of detection (Huvard et al., 2012), (Stein et al., 1995), (Cardone et al., 2006). Rapid advent of next generation sequencing technology has provided the opportunity to redesign genotyping strategies for identifying and evaluating new HNA alleles. The wide availability of population-scale genomic datasets as part of the 1000 genome (“A Global Reference for Human Genetic Variation,” 2015) and gnomAD consortium (Karczewski et al., 2020) have enabled the deeper understanding of ethnic differences in genotypes. As population-scale genome programmes accelerate in pace across the world, the paucity of a comprehensive and systematic collection of genetic variants associated with human neutrophil antigens significantly limit the understanding of population differences. To fill in this gap, we compiled a comprehensive collection of Human Neutrophil Antigen variants from literature, adding to the collection maintained by the International Society of Blood Transfusion - Granulocyte Immunobiology Working Party (ISBT-GIWP). A brief summary of the compiled genetic variants is provided in **Table 1**. This collection has a total of 37 genomic variants encompassing the five major HNA classes and is available at the Human Neutrophil Antigen Resource http://hunar.genomes.in.

**Table 1.** Tabulation of list of ISBT approved and other curated variants of HNA classes 1-5.

| HNA-1 : Antigens encoded by FCGR3B gene |  |  |  |  |  |  |
| --- | --- | --- | --- | --- | --- | --- |
| Chr | Pos (Hg38) | Ref | Alt | RSID | Category | Reference |
| 1 | 161629781 | T | C | rs2290834 | ISBT approved variant | - |
| 1 | 161629853 | T | C | rs147574249 | ISBT approved variant | - |
| 1 | 161629864 | G | T | rs5030738 | ISBT approved variant | - |
| 1 | 161629903 | T | C | rs448740 | ISBT approved variant | - |
| 1 | 161629983 | A | G | rs527909462 | ISBT approved variant | - |
| 1 | 161629989 | G | C | rs200688856 | ISBT approved variant | - |
| 1 | 161629726 | C | T | rs201867931 | Curated variant | (He et al., 2014) |
| 1 | 161629739 | C | T | rs202089363 | Curated variant | (He et al., 2014) |
| HNA-2 : Antigens encoded by CD177 gene |  |  |  |  |  |  |
| Chr | Pos (Hg38) | Ref | Alt | RSID | Category | Reference |
| 19 | 43361169 | A | T | rs201821720 | Curated variant | (Wu et al., 2019) |
| 19 | 43362297 | G | A | rs78718189 | Curated variant | (Wu et al., 2019) |
| 19 | 43361225 | A | T | NA | Curated variant | (Bayat et al., 2016) |
| 19 | 43361509 | G | - | NA | Curated variant | (Bayat et al., 2016) |
| 19 | 43361175 | A | C | NA | Curated variant | (Moritz et al., 2010) |
| 19 | 43362090 | G | A | NA | Curated variant | (Moritz et al., 2010) |
| 19 | 43353934 | A | T | NA | Curated variant | (Moritz et al., 2010) |
| 19 | 43353956 | G | A | NA | Curated variant | (Moritz et al., 2010) |
| 19 | 43362339 | G | A | NA | Curated variant | (Moritz et al., 2010) |
| 19 | 10602536 | G | A | rs115559926 | Curated variant | (Chen et al., 2016) |
| 19 | 10627963 | C | T | rs3087969 | Curated variant | (Chen et al., 2016) |
| 19 | 10631490 | C | T | rs147820753 | Curated variant | (Chen et al., 2016) |
| 19 | 10631494 | A | G | rs2288904 | Curated variant | (Chen et al., 2016) |
| 19 | 10632065 | G | A | rs146127348 | Curated variant | (Chen et al., 2016) |
| 19 | 10632155 | C | T | rs141667578 | Curated variant | (Chen et al., 2016) |
| 19 | 10635053 | A | G | rs35362308 | Curated variant | (Chen et al., 2016) |
| 19 | 10635247 | C | T | rs7255431 | Curated variant | (Chen et al., 2016) |
| 19 | 10643307 | G | A | rs34625417 | Curated variant | (Chen et al., 2016) |
| 19 | 10643310 | C | G | rs151043533 | Curated variant | (Chen et al., 2016) |
| 19 | 10643627 | T | C | rs116639546 | Curated variant | (Chen et al., 2016) |
| 19 | 10643724 | A | G | rs1053007 | Curated variant | (Chen et al., 2016) |
| 19 | 10644138 | G | A | rs74795234 | Curated variant | (Chen et al., 2016) |
| 19 | 10644229 | G | T | rs10948 | Curated variant | (Chen et al., 2016) |
| **HNA-3 : Antigens encoded by CLT2 gene** |  |  |  |  |  |  |
| **Chr** | **Pos (Hg38)** | **Ref** | **Alt** | **RSID** | **Category** | **Reference** |
| 19 | 10631490 | C | T | rs147820753 | ISBT approved variants | - |
| 19 | 10631494 | A | G | rs2288904 | ISBT approved variants | - |
| 19 | 10635012 | C | T | NA | Curated variant | (Flesch et al., 2011) |
| 19 | 10631666 | C | T | NA | Curated variant | (Flesch et al., 2011) |
| **HNA-4 : Antigens encoded by ITGAM gene** |  |  |  |  |  |  |
| **Chr** | **Pos (Hg38)** | **Ref** | **Alt** | **RSID** | **Category** | **Reference** |
| 16 | 31265490 | G | A | rs1143679 | ISBT approved variants | - |
| **HNA-5 : Antigens encoded by ITGAL gene** |  |  |  |  |  |  |
| **Chr** | **Pos (Hg38)** | **Ref** | **Alt** | **RSID** | **Category** | **Reference** |
| 16 | 30506720 | G | C | rs2230433 | ISBT approved variants | - |

## Methods

### Data retrieval and analysis

Human genome variation catalogue generated by the 1000 Genomes phase 3 project which precisely comprised over 2500 whole genome sequences from 5 major global populations and 26 subpopulations was used in the study (“A Global Reference for Human Genetic Variation,” 2015). Data was retrieved in variant call format (VCF) for chromosomes 1, 16 and 19. Information on allele counts, allele numbers and allele frequencies of each sub-populations were fetched from https://www.ensembl.org/. In addition, genetic variation information of global populations were also taken from gnomAD (version 3.1.2) (Karczewski et al., 2020). The retrieved HNA variant information was systematically documented in a pre-formatted template and was used for further downstream analysis. **Table 2**. provides a brief summary of global populations and subpopulations analyzed in the study.

**Table 2.** Tabulation of global populations and subpopulations analyzed in the study.

| **1000 Genomes Phase 3 dataset** |  |  |
| --- | --- | --- |
| **Population** | **Sub-population** | **Description** |
| African - AFR | ACB | African Caribbean in Barbados |
|  | ASW | African Ancestry in Southwest US |
|  | ESN | Esan in Nigeria |
|  | GWD | Gambian in Western Division, The Gambia |
|  | LWK | Luhya in Webuye, Kenya |
|  | MSL | Mende in Sierra Leone |
|  | YRI | Yoruba in Ibadan, Nigeria |
| American - AMR | CLM | Colombian in Medellin, Colombia |
|  | MXL | Mexican Ancestry in Los Angeles, California |
|  | PEL | Peruvian in Lima, Peru |
|  | PUR | Puerto Rican in Puerto Rico |
| East Asian - EAS | CDX | Chinese Dai in Xishuangbanna, China |
|  | CHB | Han Chinese in Beijing, China |
|  | CHS | Southern Han Chinese, China |
|  | JPT | Japanese in Tokyo, Japan |
|  | KHV | Kinh in Ho Chi Minh City, Vietnam |
| European - EUR | CEU | Utah residents with Northern and Western European ancestry |
|  | FIN | Finnish in Finland |
|  | GBR | British in England and Scotland |
|  | IBS | Iberian populations in Spain |
|  | TSI | Toscani in Italy |
| South Asian - SAS | BEB | Bengali in Bangladesh |
|  | GIH | Gujarati Indian in Houston, TX |
|  | ITU | Indian Telugu in UK |
|  | PJL | Punjabi in Lahore, Pakistan |
|  | STU | Sri Lankan Tamil in the UK |
| **gnomAD v3.1.2 dataset** |  |  |
|  | AZK | Ashkenazi Jewish |
|  | EU-NF | European Non-Finnish |
|  | EU-F | European Finnish |
|  | L-AM | Latino/Admixed American |
|  | EAS | East Asian |
|  | GME | Middle Eastern |
|  | AMI | Amish |
|  | A-AM | African/African American |
|  | SAS | South Asian |
|  | OTH | Others |

### Estimation of statistical significance

With the aim of identifying significantly distinct HNA variants, minor allele frequencies were compared among global sub-populations and statistical significance was observed using Fisher’s exact test with a P-value < 0.05. Variants identified as significantly distinct were further checked for their clinical relevance in transfusion and transplantation procedures.

## Results and Discussion

It is well established that neutrophil antigens play a central role in clinically significant transfusion and transplantation related complications in addition to other auto- and allo-immune conditions.Our analysis identified significant inter-population differences in the allele frequencies of the HNA variants. **Figure 1** provides a schematic overview of the differences in frequencies of HNA variants observed among various subpopulations of the 1000 Genomes project and gnomAD genomes. Interestingly, ITGAL_16:30506720G>C (rs2230433) variant, encoding the antithetical epitope of HNA-5a was found significantly distinct in all the 5 major global subpopulations with a frequency of 71.4% in South Asians, 13.4% in East Asians, 30.1% in Europeans, 42.1% in Africans and 18.9% in Americans. No allo antibodies against this form have been reported and characterized so far. In addition, ITGAM_16:31265490G>A substitution (rs1143679) which is a central variant in deciding the translation of either 61Arg or its antithetical form 61His was found to range from 0-0.15 in global subpopulations. This was found in concordance with the previous reports that states that HNA-4a allele frequencies in most global populations ranged from 0.9-1.0 (Xia et al., 2011), (Kline et al., 1986), (Sachs et al., 2004). Cases of fetal/neonatal alloimmune neutropenia induced by antibodies against HNA-4a/b and HNA-5a have also been reported (Curtis et al., 2016),(Porcelijn et al., 2011),(Mraz et al., 2016).

**Figure 1.**
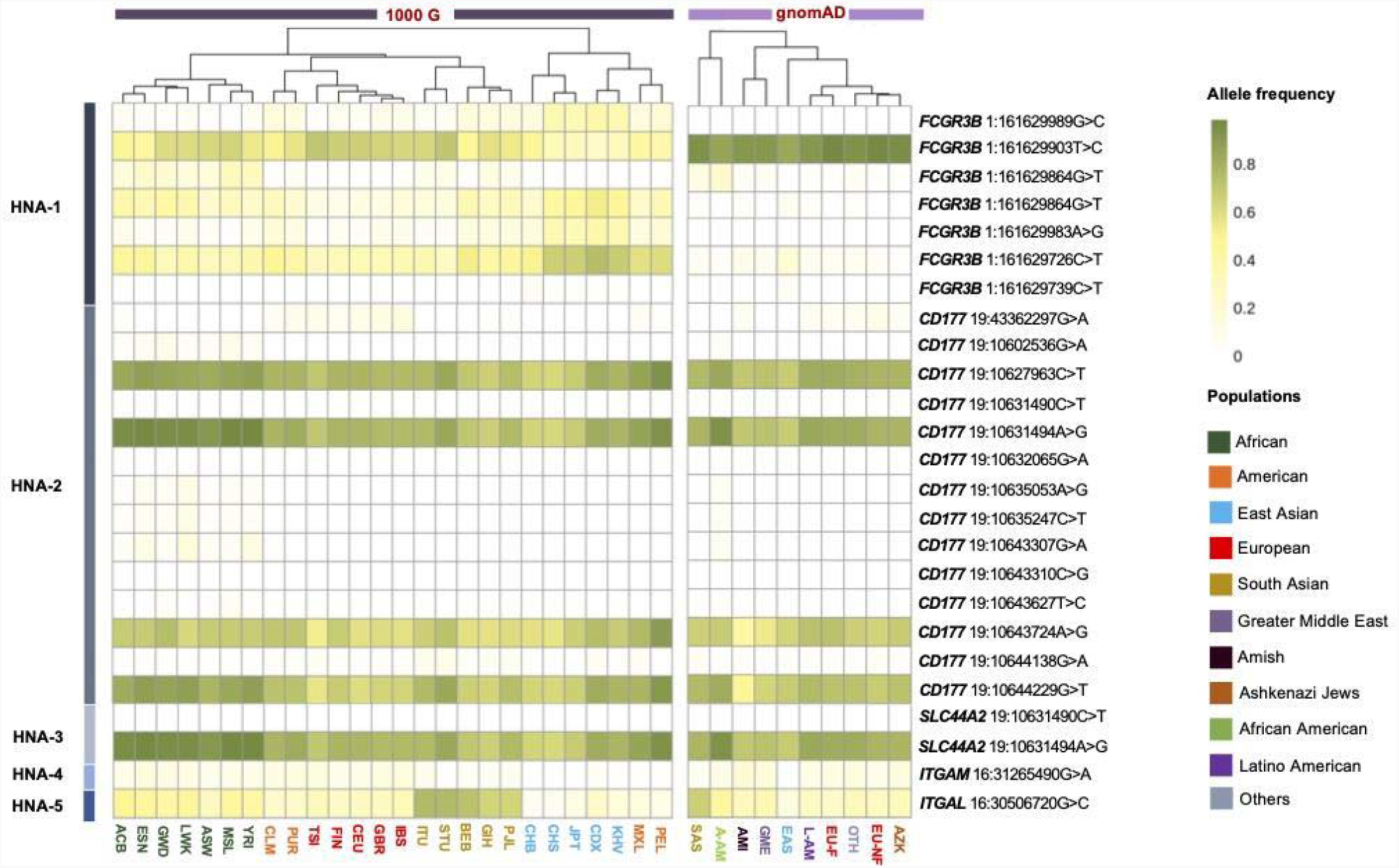
Comparison of frequencies of human neutrophil antigen variants among various subpopulations of the 1000 Genomes project and gnomAD

There also exists a handful of reports elaborating the fatal outcomes of transfusion related acute lung injury (TRALI) caused by alloantibodies against HNA-3a antigen (Chapman et al., 2009; Porretti et al., 2012),(Davoren et al., 2003). SLC44A2_19:10631494A>G substitution (rs2288904) leading to Arg152Gln marks the difference between HNA-3a and HNA-3b antigens. In compliance with previous findings, HNA-3a encoding allele frequencies were found to be 96.7% in Africans, 85.2% in Americans, 71.0% in East Asians, 76.4% in Europeans and 75.5% in South Asian populations, whereas the respective HNA-3b allele frequencies were 3.3%, 14.8%, 29.0%, 23.6% and 24.5%. The additional mutation SLC44A2_19:10631490C>T (rs147820753), which was found to impair HNA-3a epitope and abrogates the agglutination by inducing Leu151Phe amino acid change was found at a frequency ranging from 0 - 0.4% across global populations. Although concordant frequencies were observed between the two datasets used in the study, we were able to observe disparity in the frequency of FCGRB3_1:161629903T>C variant which can be possibly attributed to the fact that this variant was covered in fewer than 50% of individuals in gnomAD v3.1.2 genomes and can be a potential low-quality site.

Such efforts to estimate the genotype frequencies of neutrophil antigens among global populations provides an overview of population - specific prevalence.

## Conclusion

In conclusion, we suggest significant population differences in the HNA variants which could potentially impact clinical presentations and transfusion reactions. With the availability of genomic data from more populations which have been traditionally under-represented in global sequencing efforts, much clearer inter-population differences in HNA variant frequencies could be potentially uncovered. We sincerely hope this data could impact transfusion practices and provide much deeper insights into the genetic underpinnings of HNA related reactions and clinical presentations.

## Funding

This work was supported by The Council of Scientific and Industrial Research, India (Grant: MLP2001/GenomeApp)

## Conflicts of Interest

None declared

## Notes

### Competing Interest Statement

The authors have declared no competing interest.

